# Investigation in scanning beampattern generated by resonance cavities

**DOI:** 10.1101/2020.04.03.023663

**Authors:** Hongwang Lu, Xin Ma

**Affiliations:** School of Physics, Shandong University, Jinan 250022, Shandong, China; School of Information Science and Engineering, Qingdao 266237, Shandong, China; Shenzhen Research Institute, Shandong University, Shenzhen, China

**Keywords:** Antenna, biosonar, cavity resonance, beam forming, dipole acoustic source

## Abstract

The previous study has reported that the presence of a small flap in the Brown Long-Eared Bat (Plecotus auritus) generate useful spatial information by the frequency-driven beam scanning. However, it did not investigate the beampattern generating mechanism of the flap. In this study, according to the dipole equivalence principle, the two resonance cavities split by the flap are simplified to a dipole source with certain vibration orientation. The analytical solution of the dipole shows obvious frequency-driven sidelobe scanning in the beampattern, which matches the numerical results obtained from the pinna quite well. The results support the hypothesis that the frequency-driven sidelobe scanning is generated by the resonance in the two cavities split by the flap. This method can be applied to design a reflector antenna with small geometric structures for splitting existent resonance cavities to meet requirements of beam scanning.

---

The antenna in technical radar system can transmit the energy between the emitter/receiver and the spatial radiation. Except for that, the antenna also can obtain the azimuth angle of the target [1]. The antenna analyzing method establishes the relation between the geometrical parameters of reflector antenna and the beampatterns, so the beampatterns design can be reversely engineered to calculate the antenna geometry. Recently, with the development of the numerical computational techniques, for both the design and analysis of the reflector antenna, more complicated geometry can be designed to predict the far-field beam forming [2]. Many species of animals, especially Microchiroptera, have evolved complicated outer ear structures, which provide examples for the research on the reflector antenna as the electrical-magnetic wave and acoustic wave share similar theory. Compared to the human spatial localization, most bats of Microchiroptera can use sonar for navigation, localization, detection and preying. In the recent years, progress has been made on the effects of bats noseleaf and ear structures on the echolocation [3-12]. However, these work mostly dealt with the qualitative description of the effects of the structural change on the acoustic directivity pattern. The in-depth quantitative investigation in the mechanisms of reflection, diffraction and scattering of geometric structures and the relation between near-field and far-field were rarely mentioned. The investigation in those mechanisms can provide important theoretical foundation for the antenna design, so the mechanism that bridges the geometrical parameters and the beampattern formulation must be investigated in detail, which focuses on the effects of the structures on the near-field and far-field directivity.

The previous study showed that the maximum far-field directivity gain occurs at 32.5 kHz [5]. In Ref. [13], it was pointed out that, at 32.5 kHz, there are two resonance areas on each side of the flap, and their amplitude are very close with the phase lag pi. It was reported that the resonance around the flap is closely related to the far-field acoustic directivity in the frequency range between 30 kHz and 40 kHz. Based on the previous results, in the current study, the pinna of the Brown Long-Eared Bat (Plecotus auritus) is investigated using the field decomposition principle. According the result of field decomposition, two resonance cavities around the flap is simplified to a dipole, and the mechanism of the scanning side lobes generated by resonance cavities and dipole are analyzed by comparison of the results obtained from the numerical calculation of the real pinna and the analytical solution of the dipole.

The pinna sample of a Brown Long-Eared Bat used in this numerical experiment is obtained by the micro-CT scanning and digital processing [5,13], which is show in Fig. 1 a). As shown in Fig.1, both c) and d) is obtained by cutting the pinna model with a transverse plane marked by the frame in b). The finite element method (FEM) is employed in this study to calculate the acoustic near field. The basic computational element used is cuboid-shaped volume [5,6]. By reciprocity, the reception and radiation pattern of the ear is equivalent [14]. Hence, a point acoustic source is put in the canal opening, and then FEM is utilized to calculate the acoustic near field. The Kirchhoff integral formulation is used to calculate the acoustic far field projection. For the far field calculation, the spherical coordinate system is used. The radius of the calculated spherical surface is 10m, on which the calculated angle is −180° to 180° in azimuth and −90° to 90° in elevation both with 1° interval.

**Fig. 1.**
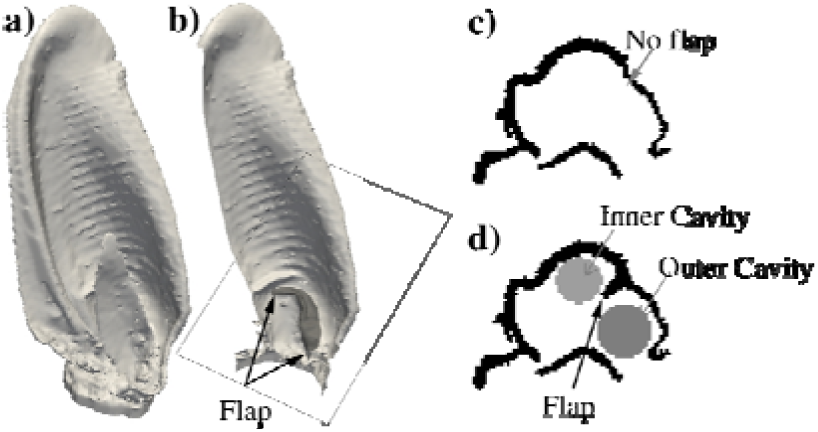
Surface rendering of the digital pinna. a) Raw pinna shape representation, and the flap is covered by tragus; b) splitting the pinna with a coronal plane to uncover the hidden flap, which is marked by arrows and dark gray area; c) transverse section of the pinna with flap removed by digital operation and d) transverse section with flap.

The acoustic near field calculated by FEM using complex numbers, for which the pressure on point can be decomposed into several vectors, as shown in Fig. 2. According to the acoustic superposition principle [14]. the overall acoustic filed is superposed by the spherical wave generated by the sound source and reflected by the baffles. Hence, the overall acoustic field can be decomposed into the summation of the wave from all sound sources and reflection. The corner reflector can be used as an example to make the concept more intuitive to be understood, which can be referred to Ref. [2,13]. One sound source is positioned in a reflector formed by two orthogonal planes, then, the acoustic near field is calculated. The total acoustic near field of corner reflector *ψ*_*C*_ can be decomposed into four parts: 1) radiation from the point sound source *ψ*_*S*_; 2) reflected sound by the vertical plate *ψ*_*V*_; 3) reflected sound by the horizontal plate *ψ*_*H*_; 4) and interaction *ψ*_*I*_. It can be written as:

**Fig. 2.**
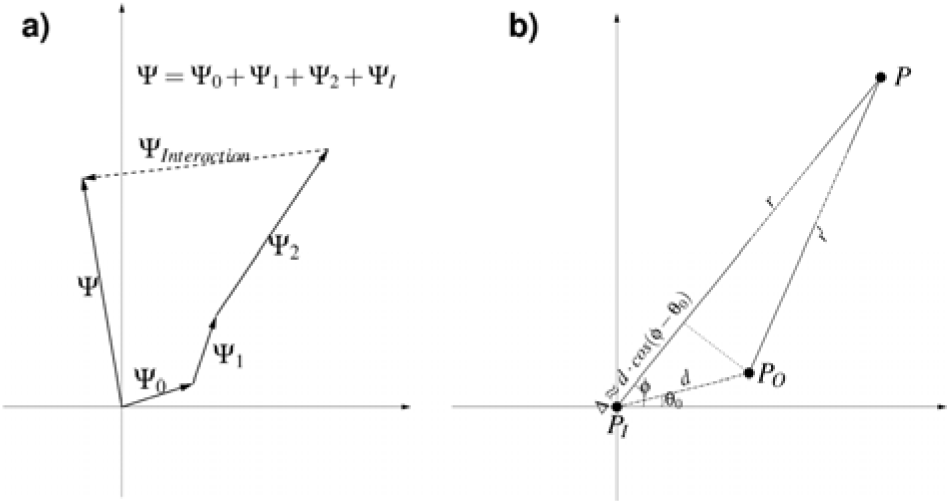
Schematic diagram of: a) resultant and decomposition of a vector, b) geometry of approximate dipole. a) A complex number is the equivalent of a 2 dimensional vector. The resultant *ψ* is the sum of *ψ*_*i*_ (where *i*=1,2,3) which can be obtained by solving the Helmholtz equation with different components flap as reflect boundary and interaction term *ψ*_*Interaction*_, which is complex difference between complete shape and other parts. b) The inner/outer cavity (*P*_*I*_, *P*_*O*_) is considered as approximate dipole with distance *d* and rotation angle *θ*_*0*_.

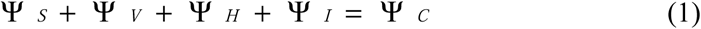

The same method can also be applied to the pinna shape, as shown in Ref [13]. For pinna shape, the total acoustic near-field can be decomposed into 4 parts: single point source (*ψ*_*S*_), reflected sound by flap (*ψ*_*F*_), reflected sound by pinna without flap (*ψ*_*N*_) and interaction (*ψ*_*I*_). Far-field directivity patterns of each part can be calculated by Kirchhoff integral formulation on the surface of near-field components. Because the wavelength is about in the same size as the flap, the sound can pass around the edge of the flap as the diffraction effect, so the flap has little effect on both near and far field. In addition, the wave of the single source is spherical wave which is isotropic, so it has no effect on directivity gain. As a result, the single source and flap items are not significant in this study. Hence, only the other three items are considered in this study.

The cavity surrounded by the pinna wall and tragus is separated into two half-open cavities by flap (see Fig. 1), which are named inner cavity and outer cavity respectively. The layer crossing the center of resonators can be picked up for calculating far-field directivity, and dipole approximation is involved to simulate the frequency-driven sidelobe scanning as the amplitude of sound pressure in both cavities increase much more than other range if resonance occurs. In the acoustic far-field (10m radius in this study), the radiation of the inner resonator can be equalized to point source *p*_*i*_, and the radiation of the outer resonator can be equalized to point source *p*_*o*_. A Cartesian coordinate system is built with *p*_*i*_ as origin, and the pressure amplitude at any point *p(r*, □) can be written as [13,

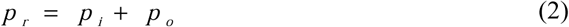

Where,

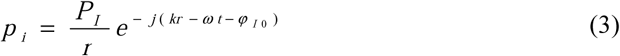

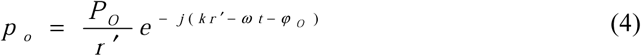

where, *P*_*I*_ and *P*_*O*_ are sound pressure amplitude of dipole, *φ*_*O*_ and *φ*_*I*_ is the initial phase, *r* and *r’* are the distance between points of dipole and the calculated point,

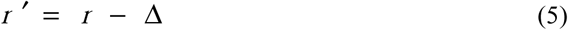

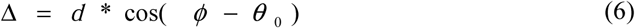

where, □ is the elevation angle, *d* is the distance of two points of dipole, *θ*_*0*_ is the angle between the dipole and the horizontal plane. The Eq. (2) can be written as

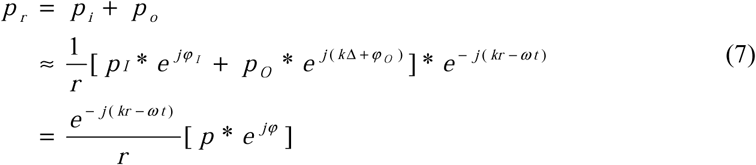

where,

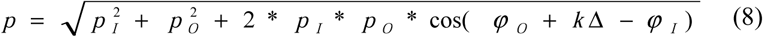

and,

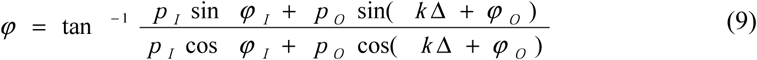

As shown in Eq. (7), when *r* is constant, *p*_*r*_ is determined by the phase difference between the dipole *φ*_*O*_*-φ*_*I*,_ wave number *k* and *d*cos(*□*-θ*_*0*_), which is the difference of sound path that the sound travels from far-field to each pole (see Fig. 2 (b)).

As shown in Fig. 1, the flap is situated in a half-opened circle, and split one cavity into two cavities. When the two cavities are in resonance, they can be simplified into a dipole (shown as Fig. 2 (b)), and this simplification is investigated in this study. The data obtained from the both cavities are analyzed. The geometrical central points of the cavities are considered as the position of the dipole, and angle between the dipole and the horizontal line is the initial oblique angle *θ*_*0*_. The amplitude (*p*_*I*_, *p*_*O*_) and phase (*φ*_*I*_ and the outer cavity *φ*_*O*_) of the dipole are calculated for each frequencies and plotted for each items (pinna without flap, flap, interaction), as shown in Fig. 3 (a), (c), (e). Meanwhile, the sidelobe magnitude in the directivity pattern is measured for all the cases, and they are plotted together with the pressure and phase ratio in Fig. 3 (b), (d), (f). In the plot, if the sidelobe maximum gain is smaller than −6 dB, it is marked as 0; if the sidelobe maximum gain is between −6 dB and −3 dB, it is marked as 0.5; if the sidelobe maximum gain is larger than −3 dB, it is marked as 0.5.

**Fig. 3.**
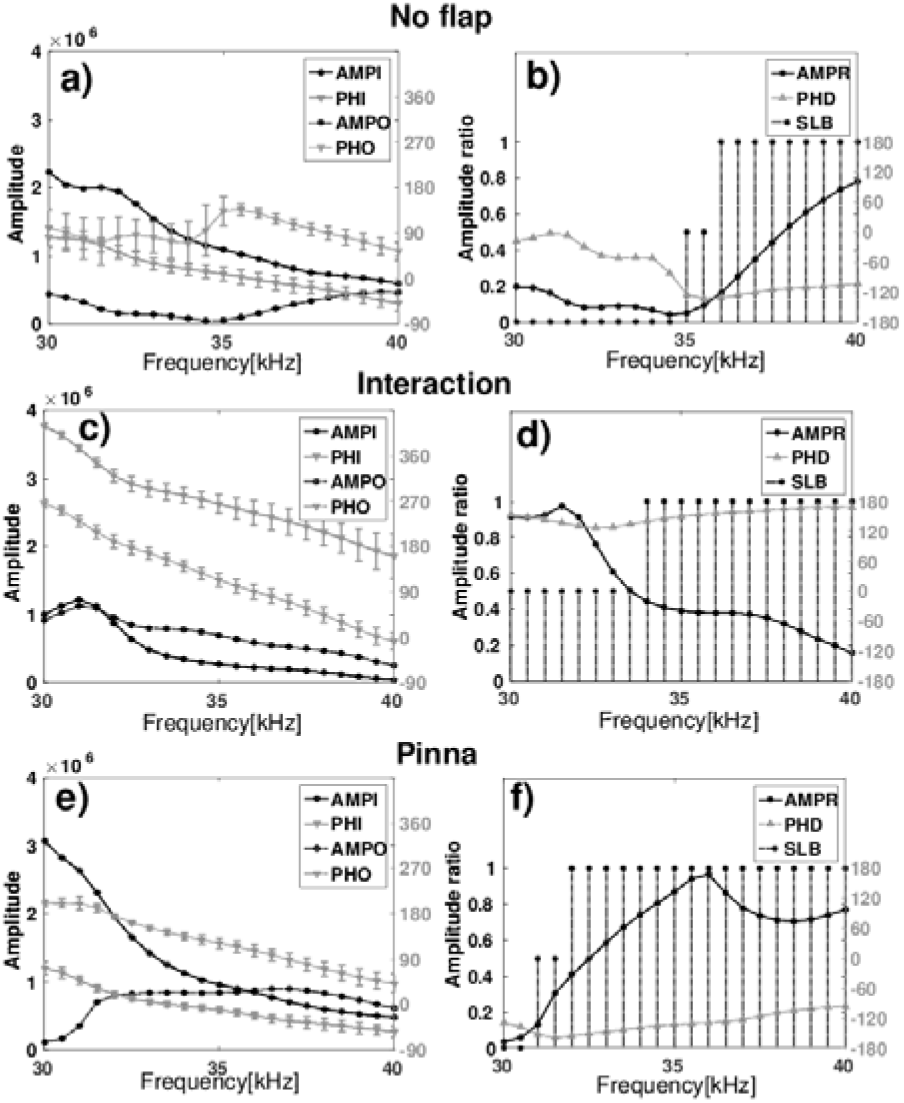
Sound pressure amplitude ratio of the approximate dipole, and phase difference (the key to generating slide lobes). a),c),e): Sound pressure amplitude (abbreviated AMP, with marker •) and phase (abbreviated PH, with gray color triangle ▾) around center of half-open cavity (FIG1.a-c) with standard deviation in inner (abbreviated I, solid line) and outer (abbreviated O, dashed line), and left Y axis is for sound pressure amplitude while right Y axis is for phase. b),d),f): Amplitude ratio calculated by minimum over maximum to measure the relative difference (AMPR, •) between “inner cavity” and “outer cavity” (left Y axis). Phase difference (PHD,Δ, right Y axis) is the difference between phase in center of inner and outer cavity, which is calculated by the linear subtraction between the inner averaged phase and outer averaged phase. Stem drawing is involved to measure the sidelobe(share the left Y axis). Let SLB=0 if there is no sidelobe more than −6dB, and SLB=0.5 if sidelobe is more than −6dB but less than −3dB, as well as SLB=1 if sidelobe is more than −3dB.

The statistical results of sound pressure and phase in near-filed around the flap in Fig. 33 show that the generation of the sidelobes is related to the out-of-phase resonance of the two cavities. It can be seen from Fig. 3 that when the pressure amplitude of the two cavities are close (ratio ≈ 1), and when they vibrate out-of-phase, large sidelobe can be generated. The frequency-driven sidelobe scanning is related to sound pressure and phase difference of resonance cavities, which can be explained by the interference of waves generated by the simplified dipole. The reinforcement is where two waves arrived in with the same amplitude and phase, while cancellation is where two waves arrive with the same amplitude but out of phase. If dipole is in phase with each other, the mainlobe of beampattern is always in the direction on the perpendicular bisector of dipole, and meanwhile it is more complicated if the dipole is out of phase. According to Eq (7), the direction of mainlobe depends on P_I_, P_O_, *φ*_*I*_, *φ*_*O*_, *k, d* and *θ*_*0*_. The directions of mainlobe and sidelobe are only dominated by *k* if other parameters are constant or changed slowly with frequency scanning, which is observed in this numerical experiment (see Fig. 3).

After the investigation in the cavities sources, the simplification of the cavities into the dipole is investigated. The averaged pressure amplitude of the two cavities *p*_*I*_ and *p*_*O*_ are taken into the dipole equation, the far-field directivity can be obtained (see Fig.4 (a)). Because the resonance cavity is above the sound emitting source (ear canal opening), so the vibration of the equalized dipole is along the Z-axis direction and not isotropic. Hence, when calculating the sound pressure on the far-field sphere, an obliquity factor should be multiplied with calculated pressure amplitude *p*_*r*_, which is *sin*□ (□ is the elevation angle). The calculated far-field directivity pattern are along the central azimuth position with the elevation angle between 0°-90° (equator to north pole) are plotted for both the simplified dipole and the pinna, as shown in Fig. 4. It can be observed that for all the decomposed sources and both models, the direction of the sidelobe changes with the frequency modulation. For the pinna without flap and the raw pinna, both models show only mainlobe at low frequencies, and frequency-driven sidelobe scanning at high frequencies, meanwhile the sidelobe enhances with frequency increasing for both models. For the interaction, there is not obvious alignment in far-field directivity for both models. The simplified dipole model only has one mainlobe as well as frequncy-driven mainlobe scanning, while the pinna numerical results contained both mainlobe and sidelobe scanning.

**Fig. 4.**
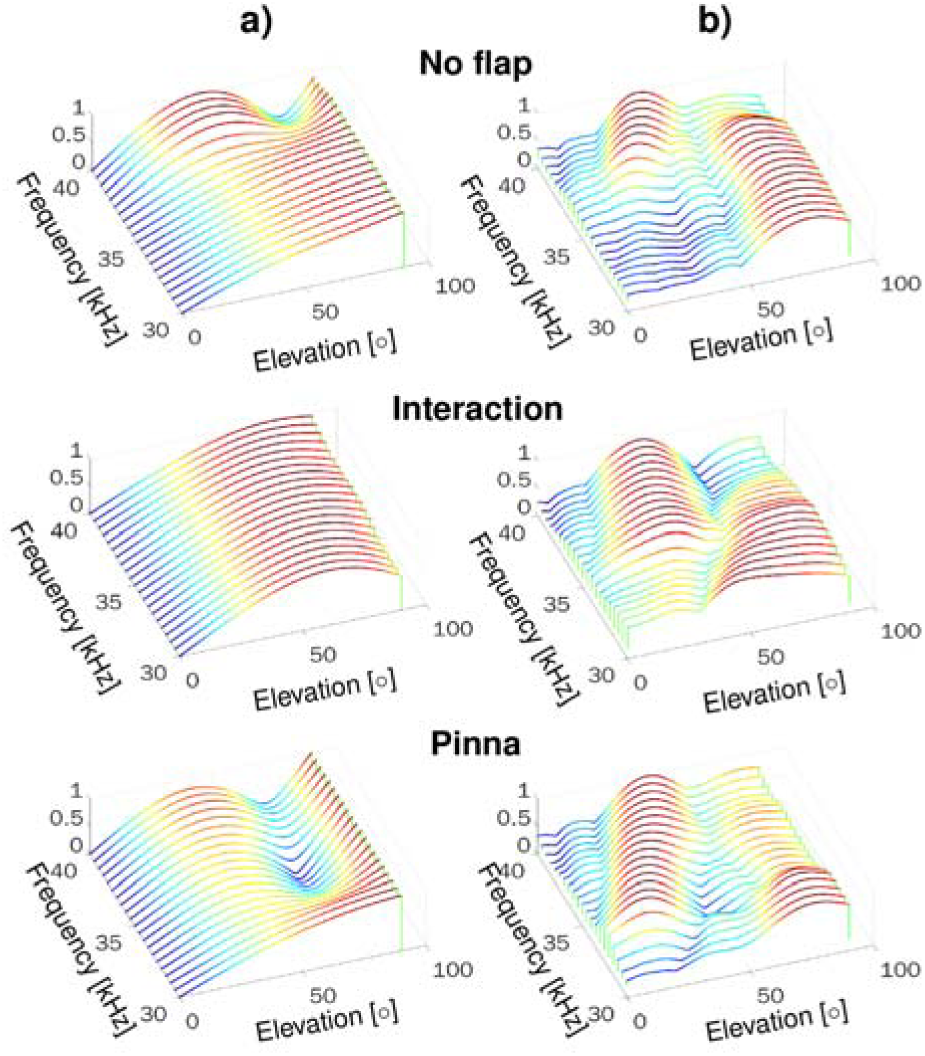
Directivities of a) simplified dipole and b) decomposed numerical model, in a fixed azimuth angle determined by direction of the simplified dipole. The X axis is elevation (0°-90° with interval 1°), the Y axis is frequency (30-40 kHz with interval 0.5 kHz) and Z (height) is normalized directivity.

In summary, the current study investigated the effects of the flap on the frequency-driven scanning. It is found that the far-field results of the dipole model match the cavity resonance quite well. Hence, it can be concluded that the far-field directivity can be manipulated by changing the resonance mode near the sound source. In addition, the current results can also be used to analyze the relation between the near-field vibration of the antenna with resonance cavities and far-field beam forming, thus inspiring the design of the reflector antenna with fixed-phase-lag resonance cavities that can perform frequency-driven sidelobe scanning. The current findings can be applied to design a reflector antenna with small geometric structures for splitting existent resonance cavities to meet requirements of beam scanning.

However, the results calculated from the decomposed interaction for the simplified model and the numerical model, which is assumed to be caused by the approximation using obliquity factor. In this study, the resonance of both cavities is approximated to be along the Z-axis, in fact, the resonance in the cavities has more complicated directions, which requires more detailed investigation in later study. Furthermore, the resonance frequency of the cavities is related to the geometrical size and the wave length of the cavities. Hence, more study on the effects of the geometric structures on the resonance amplitude and phase is needed to be done later.

## Acknowledgments

This work was supported by the Shenzhen Science and Technology Research and Development Funds (Grants No. JCYJ20170818104011781) the Key Research and Development Program of Shandong Province (Grants No. 2017GGX10113).

